# Nutritional Immunity and Antibiotic Drug Treatments Influence Microbial Composition but Fail to Eliminate Urethral Catheter Biofilms in Recurrently Catheterized Patients

**DOI:** 10.1101/637819

**Authors:** Yanbao Yu, Harinder Singh, Tamara Tsitrin, Keehwan Kwon, Shiferaw Bekele, Rodrigo V. Eguez, Yi-Han Lin, Patricia Sikorski, Kelvin J. Moncera, Manolito G. Torralba, Lisa Morrow, Randall Wolcott, Karen E. Nelson, Rembert Pieper

## Abstract

Polymicrobial biofilms that form on indwelling urethral catheters used by neurogenic bladder patients are known to recur following catheter replacements. Uropathogens dominate in catheter biofilms (CBs), grow and disperse as multi-cellular aggregates. Their microbial complexity, the characteristics of host immune responses and the molecular crosstalk in this ecosystem are incompletely understood. By surveying eight patients over up to six months with meta-omics analysis methods, we shed new light on the longitudinal microbial dynamics in CBs and the microbial-host crosstalk. There was evidence of chronic innate immune responses in all patients. Pathogens dominated the microbial contents. *Proteus mirabilis* often out-competed other species in cases of salt encrustation of catheters. The examination of proteomes in CBs and associated urinary pellets revealed many abundant bacterial systems for transition metal ion (TMI) acquisition. TMIs are sequestered by effector proteins released by activated neutrophils and urothelial cells, such as lactotransferrin and calgranulins, which were abundant in the host proteomes. We identified positive quantitative correlations among systems responsible for siderophore biosynthesis, TMI/siderophore uptake and TMI cellular import in bacterial species, suggesting competition for TMIs to support their metabolism and growth in CBs. *Enterococcus faecalis* was prevalent as a cohabitant of CBs and expressed three lipoproteins with apparent TMI acquisition functions. Fastidious anaerobic bacteria such as *Veillonella*, *Actinobaculum*, and *Bifidobacterium* grew in CB communities that appeared to be oxygen starved. Finally, antibiotic drug treatments were shown to influence microbial composition of CBs but failed to prevent re-colonization of urethral catheters with persisting and/or drug-resistant newly emerging pathogens.

## Introduction

Urethral catheter-associated urinary tract infection (CAUTI) is the most common type of complicated UTI. CAUTIs have a higher risk of recurrence, pyelonephritis and bacteremia than uncomplicated UTIs in nosocomial environments (1–3). Asymptomatic cases are usually diagnosed as catheter-associated asymptomatic bacteriuria (CAASB). The use of nearly 100 million urethral catheters per year worldwide, the 3% to 10% incidence of bacteriuria over 24 hours following patient catheterization and an average bladder catheter insertion time of 72 h (2) suggest an estimated 9 to 27 million CAUTI cases per year globally. Among the most common causes are *Escherichia coli*, *Klebsiella pneumoniae*, *Pseudomonas aeruginosa*, *Proteus mirabilis*, *Enterococcus* and *Candida spp.* (1, 3, 4). Indwelling Foley catheters are often used by patients with anatomical urinary tract abnormalities and neurogenic bladder syndrome and retained in the urinary tract for one week or longer. Microbial colonization is difficult to avoid even when catheters are regularly replaced, and antibiotic drug treatments are administered. Bacteria adapted to form biofilms (*e.g.*, *Enterococcus faecalis*, *P. aeruginosa*, and *E. coli*) and those that degrade urea and use ammonia as a nitrogen source (*e.g.*, *Proteus* and *Providencia spp.*) dominate microbial communities that form on catheter surfaces (3, 5, 6). Urea degradation alkalinizes the pH of urine and triggers the precipitation of phosphate salt crystals in this milieu, thus increasing the risk of luminal occlusion and complications such as urinary stones and kidney infections (3). Unless specific risk factors exist (*e.g.*, a compromised immune system or pregnancy), clinical guidelines do not recommend the use of antibiotics for CAASB (7). Of major concern are the genetically acquired and innate resistances of CAASB-associated bacteria against several classes of antibiotic drugs. Most of them belong to the ESKAPE group of pathogens (8). Understanding the mechanisms that drive microbial cohabitation and competition in urethral catheter biofilms (CBs) may lead to new approaches to prevent or disrupt their formation.

The pathogenesis of UTI and CAUTI has been studied extensively in the context of *E. coli* and *P. mirabilis* (4, 9–12). The innate immune system has a critical role in the recognition of and defense against invading pathogens. Their surface molecular patterns are recognized by urothelial cell effectors such as immunoglobulin A and lipopolysaccharide-binding protein. The presentation to Toll-like receptors (TLRs) triggers cytokine release and signaling events that result in leukocyte infiltration. Neutrophils are the main type of immune cells attacking and phagocytosing microbial intruders (13). Pathogen clearance results from the activities of neutrophil granular effector proteins and reactive oxygen species (ROS) as well as extracellular trapping (14, 15). Investigations of *E. coli* have implicated the neutrophil cyclooxygenase-2 in the pathogenesis of recurrent UTI (16). Recurrence is influenced by host susceptibility to and the urovirulence of strains that colonize the human intestinal tract (17).

Type I fimbriae, which are expressed by many Gram-negative bacteria, are thought to initiate mucosal colonization by binding to mannoslyated uroplakins, glycoproteins that coat the surface of urothelial umbrella cells (18). *Proteus mirabilis* produces several types of fimbriae of which the best characterized ones are the MR/P, UCF and PMF fimbriae (11). This species is the major cause of encrusted CBs where struvite and apatite minerals are deposited on catheter surfaces, triggered by urinary pH increase (3, 9, 11). Encrustation refers to the deposition of these minerals on the catheter surface along with the bacteria that thrive in this milieu. Bacterial cells in such biofilms can disperse and recolonize unoccupied catheter surfaces via swarming and adherence, processes that mediate ascendance to the kidneys and enhanced risk of pyelonephritis and urosepsis (3). The mechanisms that control microbial CB dynamics over time are complex and implicate the availability of nutrients such as carbohydrates, nitrogen, and transition metal ions (TMIs). Iron and zinc are sequestered by the innate immune system during infections (9, 19, 20). Non-encrusted biofilms have been linked to the deposition of host proteins such as fibrinogen on catheter surfaces to which bacteria adhere (3, 4). Some biofilms have a mucoid consistency due to production of extracellular polysaccharides that encapsulate bacterial cells and impede their killing by phagocytic cells. They have also been linked to renal complications by blocking urine flow (3).

Culture-based methods have shown that CBs typically harbor more than a single microbial organism (5, 6, 21). Culture-independent metagenomic surveys have identified fastidious microbes including genera such as *Actinomyces*, *Stenotrophomonas*, *Corynebacterium*, and *Finegoldia spp*. in CBs (22, 23). While these findings indicate higher microbial diversity and the ability of strictly anaerobic bacteria to colonize catheter surfaces, to what extent fastidious microbes compete with typical uropathogens and cause inflammation in the host is unclear. A few studies have analyzed microbial profiles in urine sediments or CBs pertaining to long-term catheterization of patients. One study reported high prevalence of *E. coli*, *P. mirabilis*, and *Providencia stuartii* strains in the context of persistent bacteriuria in patients catheterized over four or more weeks (24). In a study examining 4,500 urine samples from repeatedly catheterized spinal cord-injured patients, the incidences of UTI with one and two or more identified bacterial species were 45% and 15%, respectively (25). A third study revealed polymicrobial colonization by two to four common uropathogens in 20 patients (26). Notably, antibiotic treatments altered the composition of CBs and failed to clear pathogens from the patients’ urinary tracts (26).

Bladder catheterization itself was reported to cause sterile inflammation in a murine model, with CD^45+^ neutrophils as the main infiltrating immune cells. Using this model, *E. coli* and *E. faecalis* infections were shown to cause urothelial barrier disruption and further immune cell infiltration (27). Pyuria is known to occur in patients with indwelling catheters independent of the symptomology (7). Our recent work revealed neutrophil and complement system activation at similar levels in patients diagnosed with CAASB and CAUTI after catheterization at a single timepoint (28). Here, we publish the first comprehensive analyses of catheter biofilms associated with recurrent catheterization of patients. We used 16S rRNA taxonomic and proteomic analyses, and further validated results of the host-microbial crosstalk with biochemical and microbial culture methods.

## Results

### Patient cohort, antibiotic treatments and phenotypic observations

We enrolled two female and seven male human subjects of either hispanic or caucasian ethnicities. All of them had spinal cord injuries and suffered from neurogenic bladder syndrome. Comorbidities were chronically infected wounds. During the patient visits for wound treatment, indwelling bladder catheters were usually replaced. All patients received topical wound treatments with antibiotic drugs. Systemic antibiotic treatments over a limited time pertained to three patients while this study was conducted. The drug treatment regimens and other medical data are provided in Table S1 (Suppl. Data). Patient P6 was diagnosed with a renal infection a month after specimen collections ended, probably via ascension of catheter-associated pathogens to the kidneys. UTI symptoms were not reported by patients while the study was performed, which is consistent with diagnoses of CAASB. Catheters were replaced in 1- to 3-week intervals to reduce risk of CAUTI and renal complications. We analyzed urine sediments collected as centrifugal pellets from catheter collection bags and biofilms extracted from external and internal catheter surfaces. The terms used for these specimens are UP and CB, respectively, and are from here on. They were collected longitudinally from patients over 2 to 6 months, ranging from 4 to 15 timepoints. Some timepoints were represented by only a CB, a UP or both types of samples. Catheter encrustation was observed for several samples from P4, P5, P6, and P7 (Table S1, Suppl. Data). The variation in CB biomasses among the patients is displayed in Fig. 1. P6, the patient suffering a renal infection, revealed the highest average CB biomass. Four patients (P2, P4, P7, and P8) revealed only moderate variances in biomasses.

**Fig. 1.**
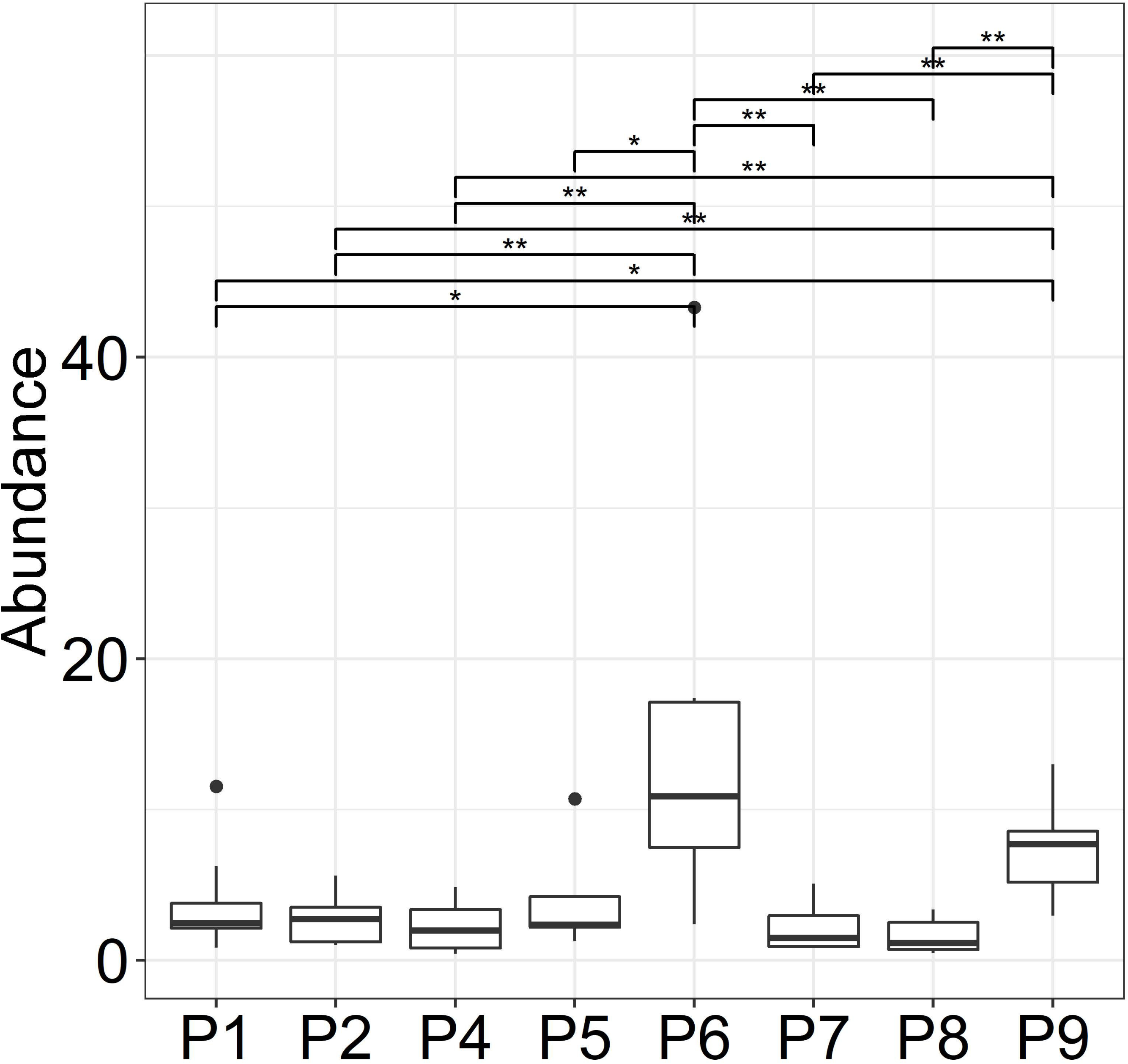
Variances in CB biomasses presented separately for each patient in box plots. We use the same patient identifiers (P1 through P9) in the main text, other Figures, and Suppl. Datasets. P3 was not included because only two data points were available. (m) male, (f) female. Biomass (abundance) pertains to wet pellet weights in g x 10^−2^ for ~ 1.5-inch catheter pieces. The CB extracts were washed in PBS, thus dissolving salt crystals that did not contribute to pellet weights. The horizontal bars depict statistically significant weight differences comparing datasets from individual patients. Significance levels are coded ***=0.001, **=0.01, and *=0.05, using the Wilcoxon rank sum test.

### Metagenomic data suggest polymicrobial colonization of catheters recurring in individual patients

Compared to microbial cultures, 16S rRNA sequencing is a less biased method to determine the taxonomic composition of a microbial community. UP and CB samples derived from 8 patients were subjected to 16S rRNA gene analysis. The data suggested the presence of 1 to 15 distinct bacterial genera in 112 specimens (Dataset S2, Suppl. Data). Species-level resolution is not achieved by sequencing the V1-V3 region of 16S rRNA. Forty genera with operational taxonomic unit (OTU) abundances greater than 0.07% (arithmetic mean) were identified. Resolution at the genus level only allows inferences of the presence of bacteria causing UTI and the urogenital microbiome. Most prevalent were Enterobacteriaceae family members and *Enterococcus*. Less prevalent were *Staphylococcus* and *Aerococcus*. Among fastidious organisms, we identified genera belonging to Actinobacteria, Bacteroides and Fusobacteria. The same genera were often repeatedly identified in samples from a distinct patient, suggesting the biological recurrence of the bacteria in sequentially replaced catheters. There was evidence of abrupt changes in the taxonomic composition for some cases that we elaborate on in a later section. Among the genera rarely associated with pathogenesis in the urinary tract were *Bordetella*, *Globicatella* and *Haemophilus*. As shown in Fig. 2, the bacterial genus diversity was moderately lower in salt-encrusted biofilms (P4, P6, and P7) compared to non-encrusted biofilms (P1, P2, and P8). P5 datasets were split into these groups as salt crystals were observed only for three early collection timepoints. This data supports the notion that phosphate salt deposition on catheter surfaces favors growth of the urease-producing *Proteus/Providencia* group of bacteria (29). Quantitative information for OTUs from 16S rRNA sequencing data is of limited value due to differences in the 16S rRNA gene amplification efficiencies (30). 16S rDNA taxonomic assignments were also useful to customize databases for proteomic analysis.

**Fig. 2.**
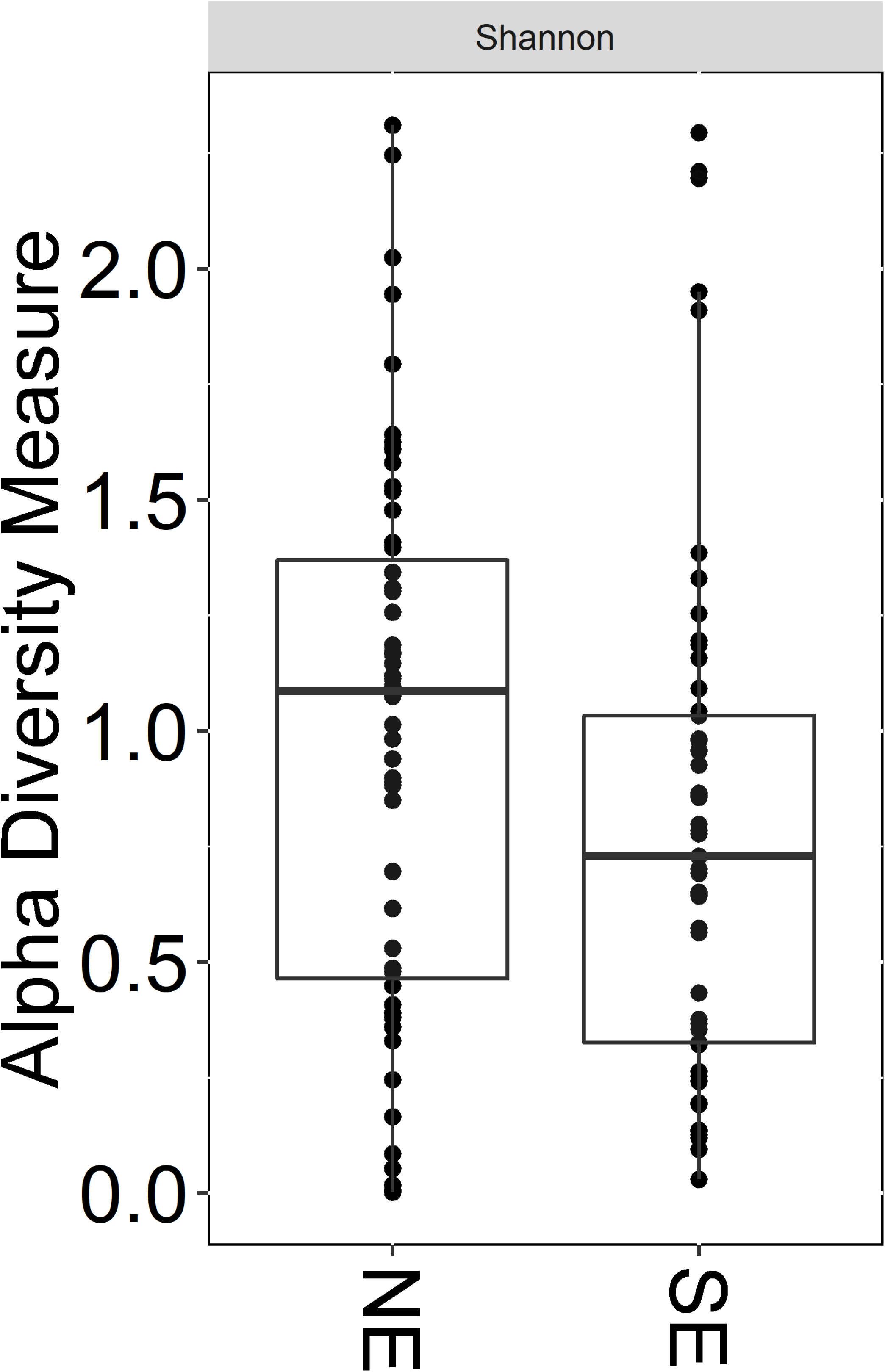
Bacterial alpha-diversity using calculations based on the Shannon index. The Shannon index accounts for abundance and evenness of OTUs. NE and SE: non-encrusted and salt-encrusted biofilms, respectively. 16S rDNA data for UP and CB samples derived from the same timepoint were separate entries. NE and SE diversities were statistically different based on a Wilcoxon rank sum test (P-value < 0.05).

### Metaproteomic data reveal dominance of P. mirabilis strains in salt-encrusted CBs and the ability of fastidious bacteria to persist in longitudinally profiled CBs

Metaproteomic database searches were performed with iterative adaptations to the composition of protein sequence databases (details in Methods section) guided by 16S rRNA results. This process allowed us to: 1. identify the correct microbial species from a given genus, most relevant in cases where more than a single species from a genus is known to colonize the human urinary tract (e.g., *Proteus, Klebsiella, Citrobacter, Enterobacter, Enterococcus, Staphylococcus, Aerococcus, Actinobaculum, Bifidobacterium*, and *Candida*); 2. select several species of the Enterobacteriaceae family for quantitative analyses only if evidence of their contributions to a sample was strong, thus avoiding incorrect peptide-spectral match assignments to orthologs with high sequence homology; 3. conduct species-specific pan-proteome database searches to assess whether key proteins were missed by analyzing data from only a single genotype (strain). The latter was most insightful for *E. coli* due to a high number of sequenced strains and the identification of plasmid-encoded virulence and antibiotic resistance proteins for all species. The Table S3 (Suppl. Data) contains the protein sequence databases used for iterative searches to selectively identify the species for quantitative proteomic analyses applied to all 121 samples, including 42 collection timepoint-matched UP and CB samples. P3 was excluded from further analysis due to a low number of samples. Quantified at the microbial species level, the data is displayed in longitudinal UP and CB sample series in the graphics of Fig. 3. We were confident that this data accounted for most of the microbial biomasses. Comparisons of equivalent timepoints (16S rRNA data vs. proteomics) suggest that low abundance microbial constituents present in samples according to 16S rRNA data were absent in proteomic profiles, consistent with the notion that lower protein detection limits for such species were reached.

**Fig. 3.**
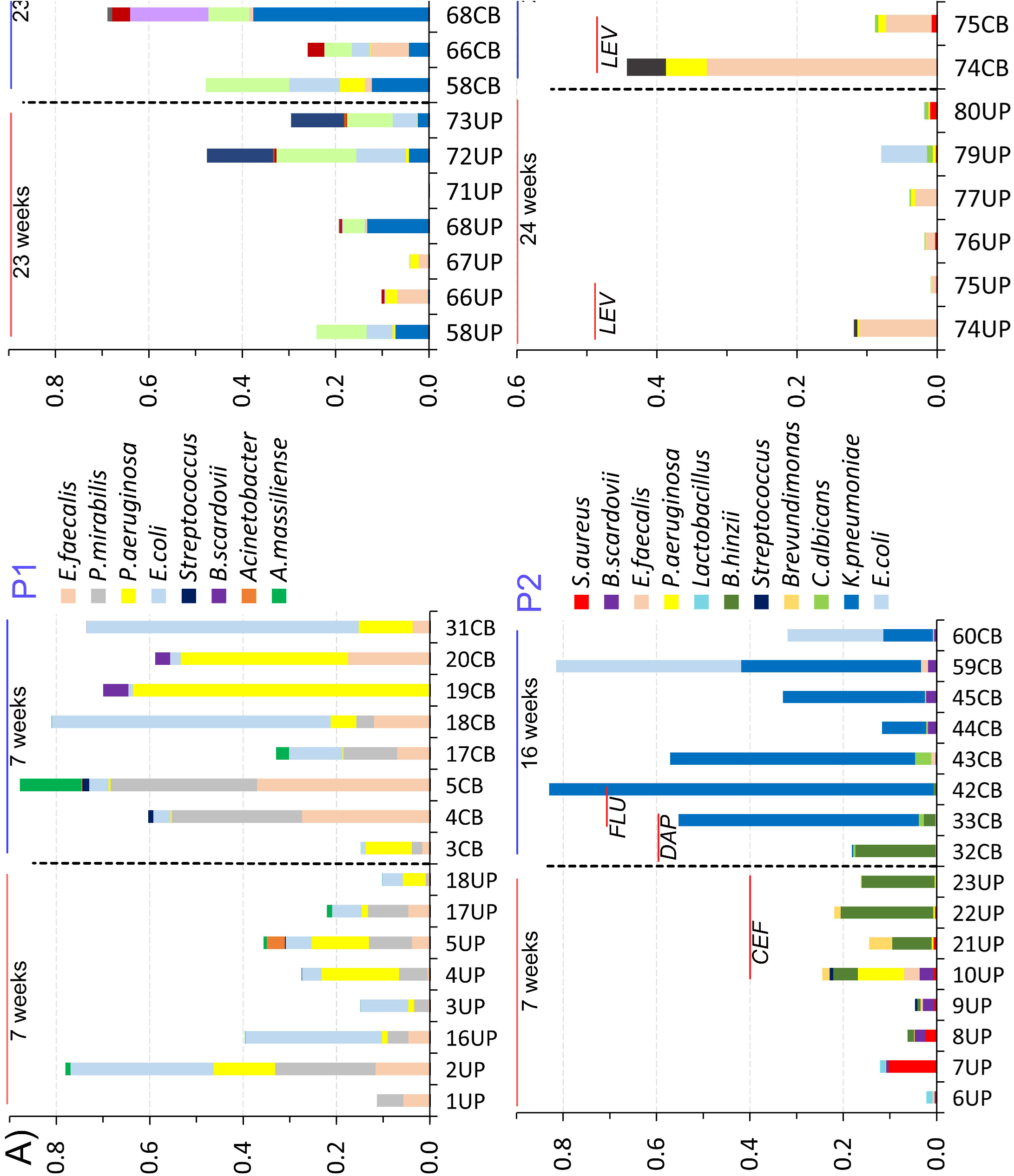
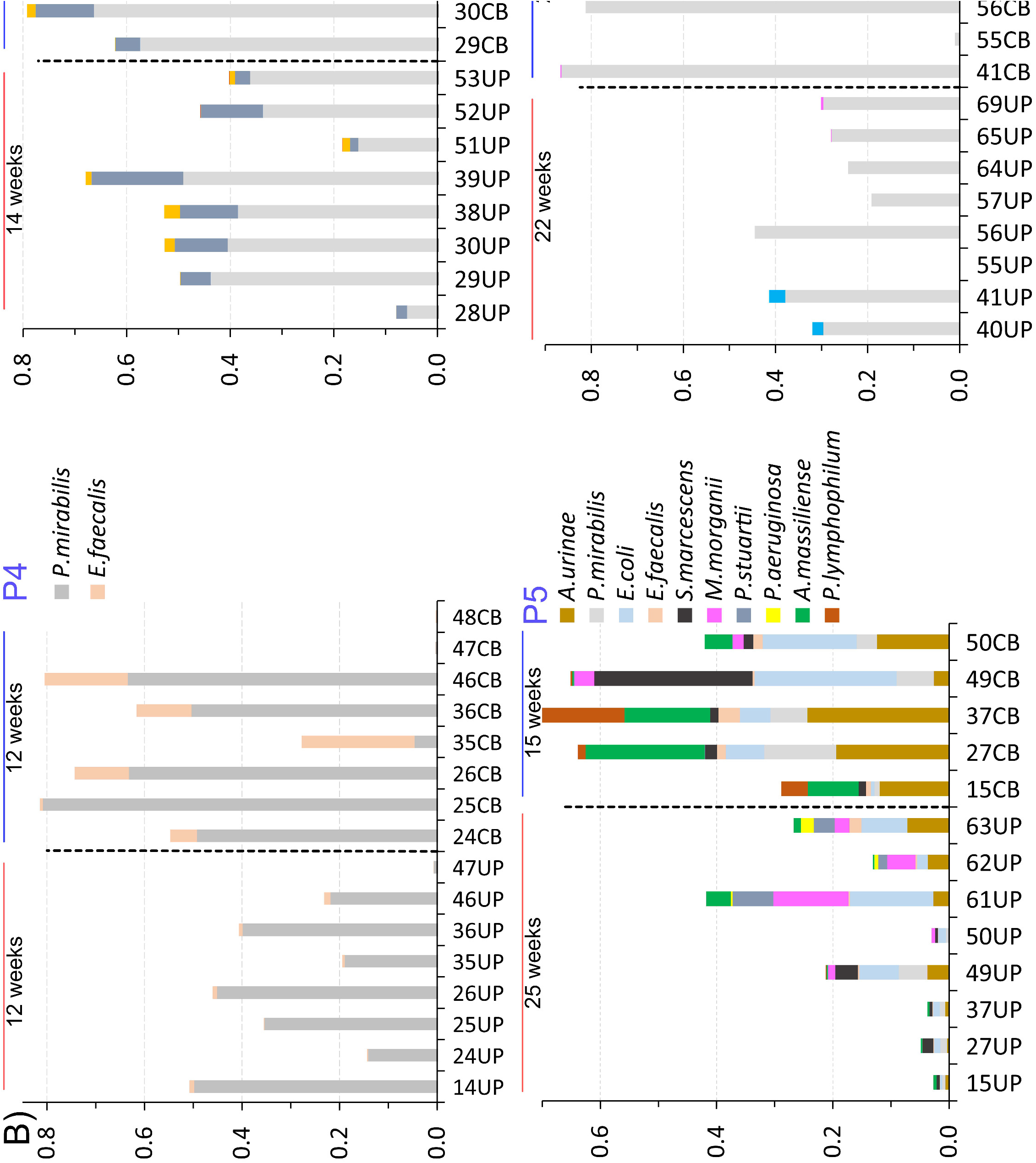
Quantitatively represented microbial species according to metaproteomic data for CB and associated UP samples. **A.** P1-P9, no evidence of salt crystals on catheter surfaces and in urine. **B.** P4-P7, evidence of salt crystals on catheters and in urine. The bars are ordered from the first to last collection time point (left to right). Collection time-matched CB and UP samples have the same number. The colored segments of bars represent the abundances of microbial species, based on the sum of their PSMs relative to the entire proteome identified from a sample. Horizontal bars at the top show the time frame of sample collection. Vertical hatched bars separate UP from CB sample sets. Inserts display the times during which antibiotics were administrated systemically (*BAC*, Bactrim; *CEF*, cefaroline-fosamil; *DAP*, daptomycin, *FLU*, fluoconazole, *LEV*, levofloxacin).

Microbial *in vitro* culture methods were used to verify species identities and recover isolates for drug susceptibility tests (DSTs). Specimens were typically stored at −80°C prior to revival of the microbial strains on media under aerobic growth conditions *in vitro*. A P5 catheter corresponding to 61UP was also preserved in liquid N_2_ after collection and grown anaerobically in rich media, As reported (31), colonies were subjected to 16S rRNA and proteomic surveys and largely agreed with the profiles derived from equivalent clinical samples. Lower abundance fastidious bacteria, such as *Prevotella*, were identified by 16S rRNA sequence analysis from both clinical samples and colony isolates. This supported the notion that microbes of low abundance in UP and CB samples were not always in a detectable range for shotgun proteomics. Using frozen specimens for colony isolation, only species that are viable during extended storage at −80°C were identified. This included *Serratia marcescens*, *K. pneumoniae*, *S. aureus*, and *E. faecalis* (Dataset S2, Suppl. Data) all of which have cell walls encapsulated with protective exopolysaccharides. In contrast, species that were quite abundant in some clinical specimens, *e.g. E. coli* and *A. urinae*, were not isolated under these conditions.

Microbial proteins quantified by peptide-spectral counting from all UP and CB samples (Dataset S4, Suppl. Data) were summed at species levels. One fungal species, *Candida albicans*, was identified in P2 and P9 datasets at low levels. Common bacterial pathogens of the urinary tract were more persistent in longitudinal profiles than rare pathogens and fastidious anaerobes (Fig. 3). *P. mirabilis* was dominant in samples from three patients with salt-encrusted CBs surveyed over 3.5 to 5.5 months (P4, P6, and P7), but less so in P5 where salt encrustation was observed for less than half of the samples. Both *P. stuartii* and *E. faecalis* persisted in two of the patient sample series. Non-encrusted catheters (P1, P2, P8, and P9) were more dynamic in microbial content of CBs over time. Most prevalent were *P. aeruginosa*, *E. coli* and *E. faecalis*. In the CBs of P2 that followed three antibiotic treatment courses, *K. pneumoniae* was dominant. Among rare pathogens, *Brevundimonas* (10UP-22UP), *Stenotrophomonas* (71CB), and *Acinetobacter* (5UP) colonized transiently, whereas *Bordetella hinzii* (10UP-33CB) and *S. marcescens* (15CB-50CB) displayed higher persistence over time. The box plots in Figure S5 (Suppl. Data) depict variances in bacterial species abundances comparing UP and CB proteomes in a patient-specific manner. No statistically significant differences in microbial composition for timepoint-matched UP and CB datasets were identified. This analysis suggests that the cycle of biofilm formation and dispersal occurs at the microbial community level. For the most part, UP samples contain those organisms that dispersed from CBs.

### Proteomic data allow assessments of strains dominance and microbial adaptation to the catheter biofilm milieu

Little is known about molecular adaptations of species to the complex microbial environment in CBs and the presence of the host’s immune cells and their effectors at the catheter-urothelial interface. Furthermore, it was of interest to determine if specific strains of a microbial species were dominant in a series of samples from a given patient. We investigated these questions patient-specifically at the proteome level. Statistically significant abundance differences were identified for 664 bacterial proteins (ANOVA tests; Dataset S6, Suppl. Data). Interpreting all data is beyond this article’s scope. We elaborate on *P. mirabilis*, *E. coli*, and *E. faecalis* because their proteomes were well-represented in datasets from several patients, with a focus on proteins relevant to bacterial energy metabolism and interactions with the host environment. HpmA was detected only in *P. mirabilis* proteomes pertaining to CBs of P7 (Fig. 4), supporting the notion that this putative hemolysin is expressed by few uropathogenic *P. mirabilis* strains. High abundance of HpmA in all P7 datasets suggests that a single *P. mirabilis* strain highly dominates the CB series of P7. A yersiniabactin-like iron/siderophore receptor (gene locus PMI2596) was abundant in the *P. mirabilis* proteomes of CB samples from P1 and P4, lower in abundance in P5 and absent in P6 and P7 samples (Fig. 4). The gene cluster for biosynthesis of yersiniabactin and expression of its receptor FyuA is considered part of the *E. coli* accessory genome. The abundances of the *E. coli* proteins FyuA and Irp1 (an enzyme encoded by one gene in this cluster) were also highly variant and most abundant in P1 and P2 datasets (Fig. 4). Some *E. coli* strains produce another siderophore, aerobactin, via the system IucA-D. Its receptor IutA (Fig. 4) and the L-lysine 6-monooxygenase IucD were differentially abundant in the *E. coli* proteomes of P1, P2, P5 and P8. Box plots displaying the variances of additional proteins involved in TMI uptake pathways among patients are included in Dataset S6 (Suppl. Data). In support of a single strain’s dominance for a species in the CB series pertaining to a distinct patient, an *E. coli* plasmid-encoded ferric iron uptake system (UTI89_P010-P017) and the Hek adhesin factor were identified in multiple samples from P1 but not in those from any other patient.

**Fig. 4.**
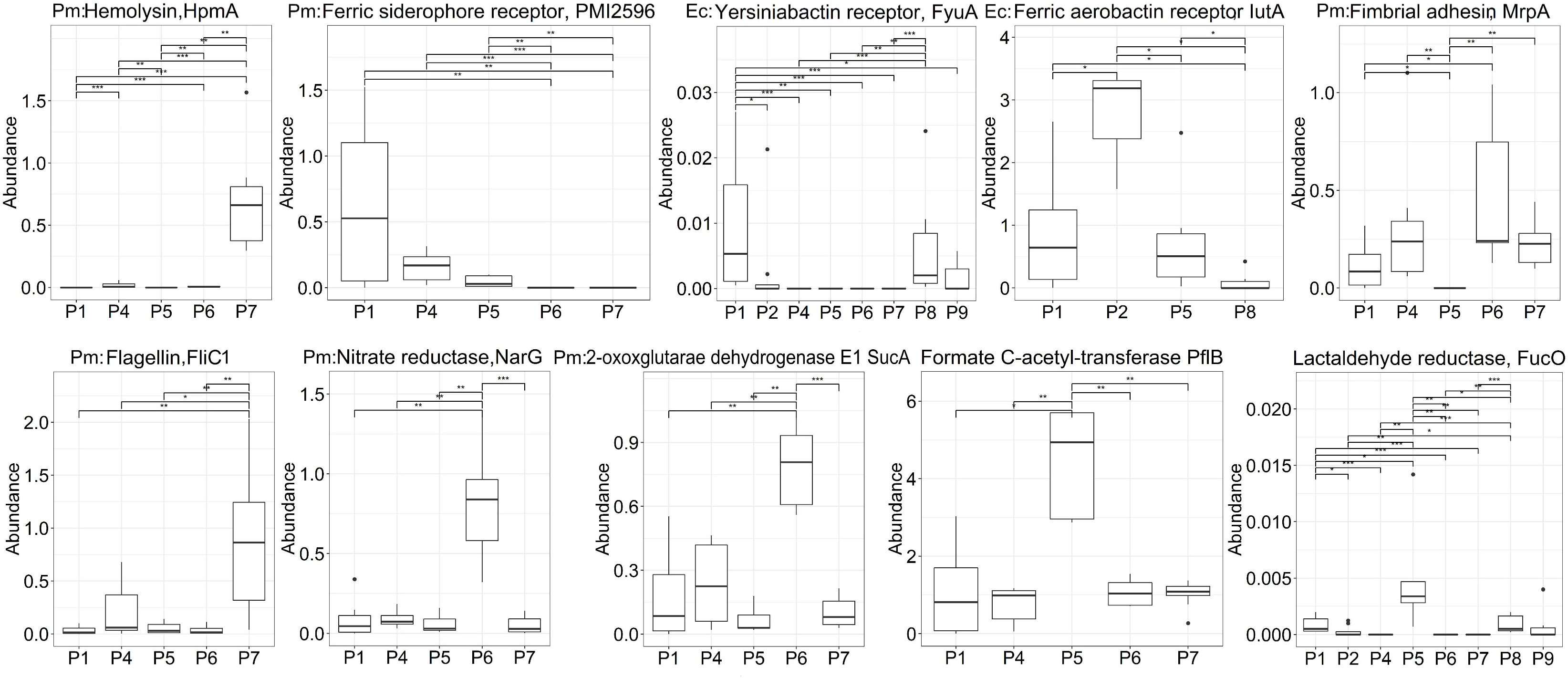
Variances in abundance for *P. mirabilis* (Pm) and *E. coli* (Ec) proteins. Proteins are listed with a short name or gene locus. Locus and conserved sequence data predict the siderophore receptor role of PMI2596. Statistical significance of protein variances is depicted by horizontal bars at the top of each box plot. The significance levels are coded as ***=0.001, **=0.01, and *=0.05. The Wilcoxon rank sum test was used to determine statistically significant differences in patient-to-patient comparisons (p-value < 0.05).

We observed statistically significant abundance differences for bacterial proteins that contribute to fitness and survival in the human host, including the *P. mirabilis* MR/P fimbriae and the flagellin FliC (Fig. 4). The mannose-resistant fimbrial protein MrpA adheres to urothelial cells and catheters. FliC enables cell swarming and likely contributes to the pathogen’s spread along catheter surfaces. MrpA was most abundant in P6, a case associated with very high CB biomasses and diagnosis of a renal infection. We identified differences in abundance for enzymes part of energy metabolism pathways in patient-specific proteome comparisons. The anaerobic respiration pathway Nar/Fdo (*P. mirabilis*), with nitrate as electron acceptor and formate as electron donor, was much more abundant in CBs from P6 than in CBs from other patients. NarG profiles are shown in Fig. 4. Abundance profiles were similar for SucA, a 2-oxoglutarate dehydrogenase subunit part of the citrate cycle. The citrate cycle produces reducing equivalents for nitrate reductase (Nar). Enzymes of the mixed acid fermentation (MAF) pathway were most abundant in CBs of P5, P7, and P8. The abundance profile of *P. mirabilis* formate C-acetyltransferase (PflB) is depicted in Fig. 4. Aldehyde-alcohol dehydrogenases (AdhE) of *P. mirabilis* and *E. coli* were also differentially abundant (Dataset S6, Suppl. Data). Fittingly, fastidious bacteria were high biomass contributors only in samples from P5 (*A. massiliense* and *Propionimicrobium lymphophilium*) and P8 (*Campylobacter curvus* and *Veillonella parvula*). A fucose degradation pathway of *E. coli* appeared to be most active in P5 samples, with statistically significant differences in abundance for FucO, FucU and FucI. Lactaldehyde reductase FucI (Fig. 4) catalyzes the terminal step of anaerobic fucose metabolism. Many proteins with functional and structural roles in mRNA and protein biosynthesis (*P. mirabilis, E. coli*, and *E. faecalis*) had statistically significant abundance differences comparing the patient groups. In summary, the differential use of proteins and multi-molecular entities important to maintain the fitness of bacteria in biofilms (mobility, adhesion, energy metabolism), comparing CB series between patients, is indicative of adaptations influenced by host and microbial environments.

### Functional correlations among molecular systems involved in TMI acquisition

We assessed whether multi-subunit systems involved in TMI acquisition, often regulated by the iron starvation-sensing transcription factor Fur, positively correlated in expression levels. The proteomes of a subset of bacteria sharing CB niches (*E. faecalis*, *A. urinae*, *P. mirabilis*, and *E. coli*) in patients P1, P4, and P5 were targeted. Abundance correlation R-values of TMI acquisition systems derived from the individual patients’ longitudinal CB timepoints were determined. There were no statistically significant correlations for P4 and P5 data. Strong positive correlations for *P. mirabilis*, *E. coli* and *E. faecalis* TMI acquisition systems, or proteins part of these systems, were observed for data from P1 (Fig. 5). These systems’ abundances did not correlate with that of Fur itself, as shown for *P. mirabilis* Fur in Fig. 5. This transcription factor activates the expression of iron acquisition systems via a mechanism not dependent on its own abundance. The correlation analyses demonstrate that microbial cohabitants jointly respond to the starvation of iron and other metal ions and express a versatile repertoire of proteins dedicated to their extracellular capture and import into the cell. The summed abundance of *P. mirabilis* ExbB and ExbD, subunits of the energy-transducing Ton system responsible for the proton motive force-dependent uptake of TMI/siderophores via TonB-dependent outer membrane receptors, positively correlated in abundance with the receptors they serve. MR/P fimbriae of *P. mirabilis* (MrpA-H) did not positively correlate in abundance with any TMI acquisition proteins. Three *E. faecalis* lipoproteins, subunits of ABC transporters predicted to bind TMIs, were expressed by the strains that cohabitated CBs of five patients: EF2076, EF0577, and EF3082. The abundances of EfaA (EF2076) varied comparing the *E. faecalis*-containing CB proteomes from five patients (Fig. 6A) and positively correlated with those of TMI acquisition systems of *P. mirabilis* and *E. coli* (Fig. 5). The data were similar for the sum of subunits of the ABC transporter (EfaABC). In a recent study, the system was found to be required for Mn^2+^ import and CAUTI pathogenesis (32). We expressed these *E. faecalis* lipoproteins recombinantly in *E. coli*, purified them, and determined protein melting profiles (T_m_). Complex formation with small molecules, *e.g.* TMIs, stabilizes a protein and increases its T_m_. We observed that purified EfaA and EF0577 had two T_m_ maxima in fluorescent dye-binding experiments. One T_m_ maximum appeared to represent an apoprotein binding a TMI as a cofactor. Denaturation eliminated this higher T_m_ peak, while renaturation in the presence of 50 µM CoCl_2_ resulted in restitution of the high T_m_ maxima (Fig. 6). Protein melting profiles for EF3082 did not display two T_m_ peaks for any experimental condition. We infer that EfaA and EF0577 bind one or more TMIs and facilitate their ABC transport-mediated uptake into the *E. faecalis* cell.

**Fig. 5.**
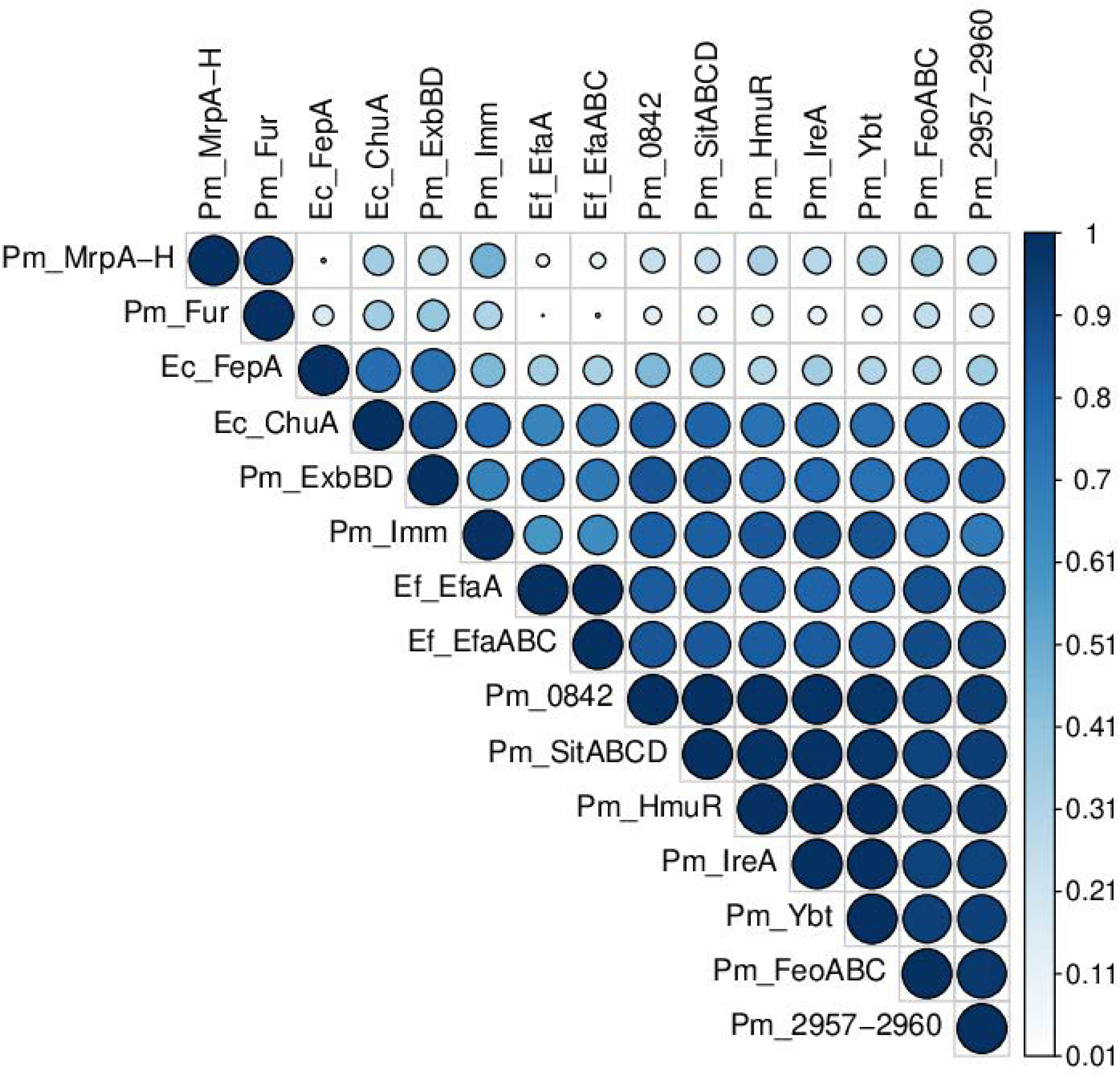
Abundance correlations of proteins and molecular systems involved in TMI acquisition. The data pertain to eight CB proteomes derived from P1. The bacterial species are Ef: *E. faecalis*; Pm: *P. mirabilis*; and Ec: *E. coli*. Correlations were determined using the Pearson correlation method. Color intensity and size of the circles are proportional to the correlation coefficients. Proteins are listed with UniProt short names or gene loci.

**Fig. 6.**
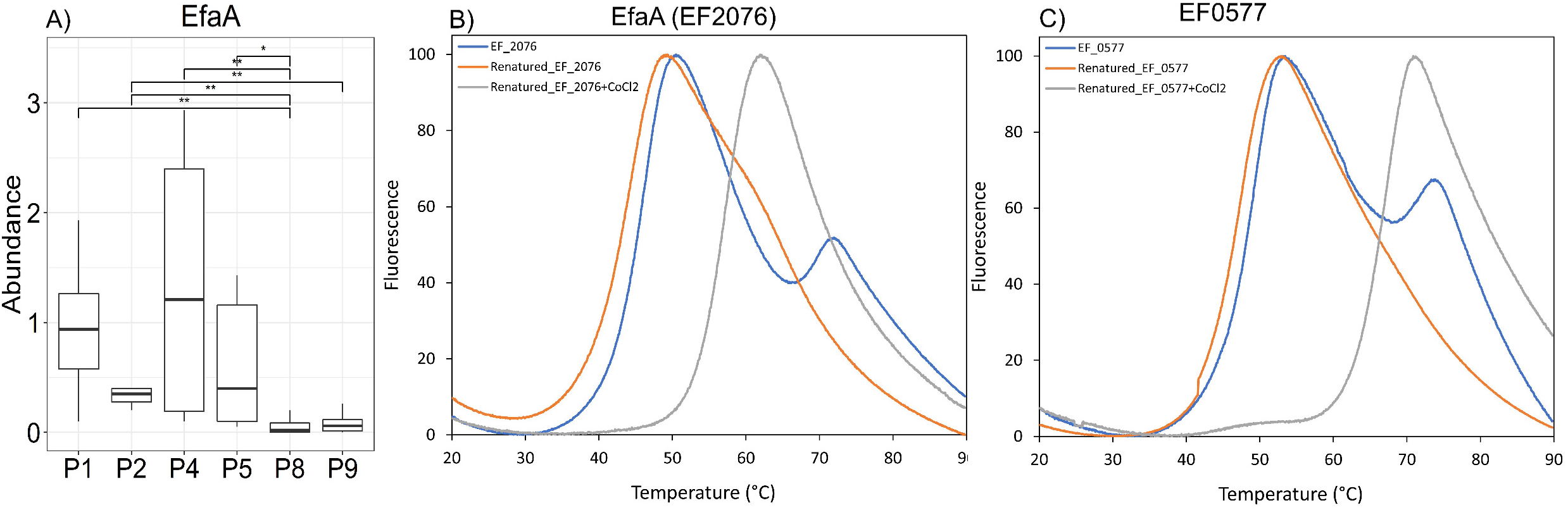
Studies on two predicted TMI-binding *E. faecalis* lipoproteins. **A.** EfaA abundance levels comparing CB data from five patients (see also legend, Fig. 4). Protein melting profiles (T_m_) for **B.** EfaA and **C.** EF0577 binding SYPRO Orange for detection using the Light Cycler 480. Recombinant proteins (12.5 μM) were equilibrated in PBS containing 1 mM DTT. T_m_ profiles are shown for purified proteins (blue), proteins denatured with 6 M guanidine-HCl and washed with 8 M Urea to remove potential bound cofactors followed by renaturation with PBS (orange) or CoCl_2_/PBS (grey) on a Ni-NTA agarose column. The concentrations of SYPRO Orange and CoCl_2_, where applicable, were 8.3X and 50 μM, respectively.

### Chronic innate immune responses result from persistent microbial colonization in all patients

The absence of CAUTI symptoms in recurrently catheterized neurogenic bladder patients does not suggest absence of innate immune responses and pyuria (7, 28). Longitudinal human proteomic profiles allowed us to assess patient- and timepoint-specific quantitative differences in the immune responses. First, we quantified relative protein abundances averaged from all 121 samples (Dataset S4, Suppl. Data). In Table 1, a subset of proteins with either a role in neutrophil- and eosinophil-mediated immune responses or as a cell biomarker is selected. Hemoglobin is a urine biomarker of tissue injury in the urethral and bladder mucosa. Hemoglobin subunits, *e.g.* HBB (Table 1), were moderately abundant in most datasets suggesting the occurrence of microhematuria in catheterized patients. High abundance of neutrophil-enriched and moderate abundance of eosinophil-enriched effector proteins (MPO, calgranulins and EPX, Table 1) suggest that both innate immune cell types infiltrated the urinary tract upon microbial colonization of the catheters. Based on a panel of cell-specific surface markers, as identified by the HCDM resource (33), granulocyte (CEACAM8) and neutrophil (CD177) markers were more abundant than B-cell, T-cell, macrophage and dendritic cell markers (Table 1). Urothelial umbrella cell surface markers were of very low abundance compared to keratins, consistent with the occurrence of squamous urothelial metaplasia and progressing epithelial cell keratinization due to chronic irritation of the patients’ urinary tracts (34). Uroplakin-2 and KRT13 are included in Table 1. Complement system proteins such as C3 were also abundant, indicative of high activity of this innate immunity branch. Uromodulin, a protein forming polymerized gel-like aggregates in urine, is abundant regardless of immune cell infiltration and therefore served to normalize protein abundance for the quantitative analyses shown in Fig. 7.

**Table 1.** Selected proteins representing different cell types and functions in innate immunity

| UniProt ID | Rk'No. | $A_{\text{Avg}}^2$ | Protein name | Functional and cell surface biomarker roles for proteins<sup>3</sup> |
| --- | --- | --- | --- | --- |
| P07911 | 1 | 3.05 | Uromodulin (UMOD) | secreted from renal tubular cells into urine |
| P02788 | 2 | 2.02 | Lactotransferrin (LTF) | abundant in activated neutrophils, iron-sequestering |
| P05164 | 4 | 1.55 | Myeloperoxidase (MPO) | abundant in activated neutrophils (azurophilic granules) |
| P13646 | 5 | 1.50 | Keratin, type I cytoskeletal 13 (KRT13) | abundant in stratified squamous epithelial cells |
| P01024 | 7 | 1.32 | Complement system component C3 | abundant effector of the complement system |
| P06702 | 10 | 0.89 | Protein S100-A9 (calgranulin) | abundant in activated neutrophils, $\text{Zn}^{2+}$ sequestering |
| P05109 | 32 | 0.45 | Protein S100-A8 (calgranulin) | abundant in activated neutrophils, Zn<sup>2+</sup> sequestering |
| P68871 | 46 | 0.33 | Hemoglobin subunit beta (HBB) | abundant in erythrocytes |
| P11678 | 51 | 0.28 | Eosinophil peroxidase (EPX) | abundant in activated eosinophils (secreted granules) |
| P80188 | 92 | 0.14 | Lipocalin-2 (LCN2) | abundant in activated neutrophils (gelatinase granules) |
| P31997 | 252 | 0.04 | Carcinoembryonic antigen-related cell adhesion molecule 8 (CEACAM8) | granulocyte cell-specific surface marker<sup>3</sup> |
| Q8N6Q3 | 421 | 0.02 | CD177 antigen (CD177) | neutrophil cell-specific granule and surface marker<sup>3</sup> |
| P20702 | 769 | 0.01 | Integrin alpha-X (ITGAX) | T-cell, B-cell, NK-cell, DC, MAC/monocyte, granulocyte cell-specific marker<sup>3,4</sup> |
| P16422 | 940 | <0.01 | Epithelial cell adhesion molecule (EPCAM) | epithelial cell specific marker<sup>3</sup> |
| O00526 | 2484 | <0.01 | Uroplakin-2 | urothelial umbrella cell-specific marker (bladder) |
| P09693 | 4898 | <0.01 | T-cell surface glycoprotein CD3 γ chain | T-cell-specific surface marker<sup>3</sup> |
| P20138 | 8191 | <0.01 | Myeloid cell surface antigen CD33 | DC, granulocyte, MAC/monocyte, stem cell marker<sup>3,4</sup> |
| P15391 | 8238 | <0.01 | B-lymphocyte antigen CD19 | B cell, DC and stem cell surface marker<sup>3,4</sup> |

<sup>1</sup>Normalized abundance rank and <sup>2</sup>Normalized relative protein quantity average from all UP and CB datasets (the PSMs matched to a protein divided by the sum of all human PSMs); <sup>3</sup>Cell differentiation marker according to (33); <sup>4</sup>abbr.: DC, dendritic cells; MAC: macrophages; NK, natural killer cells;

**Fig. 7.**
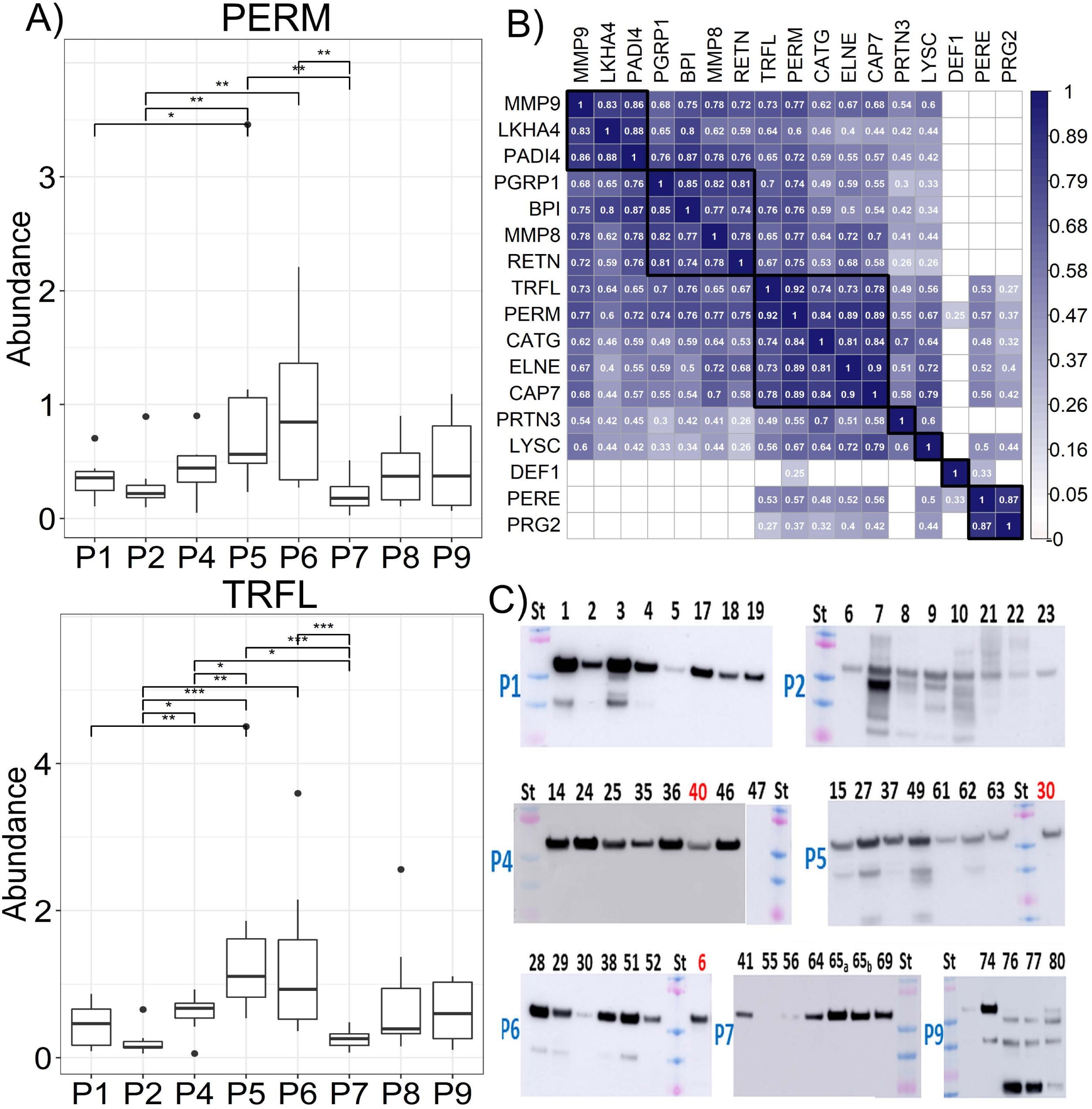
Neutrophil and eosinophil infiltration in response to persistent catheter colonization. **A.** Variation in MPO (PERM) and LTF (TRFL) abundance profiles for eight patients. Wilcoxon rank sum tests were used to assess variance. Statistically significant differences pertained to patient-to-patient comparisons (p-value < 0.05). **B.** Correlation analyses of neutrophil and eosinophil proteins were determined using the Pearson correlation method. Coloration is proportional to the correlation coefficients, and blank squares represent non-significant correlation (P-value < 0.05). Distinct clusters are boxed: this includes eosinophil peroxidase (PERE) and bone marrow proteoglycan PRG2; MPO and LTF clustered with cathepsin G (CATG), elastase (ELNE), and azurocidin (CAP7). **C.** MPO western blot data for samples derived from seven patients. Lane identifiers match UP sample identifiers also used in Fig. 3. Lane identifiers 40 (in P4 gel), 30 (in P5 gel) and 6 (in P6 gel) are not part of these patients’ sample series. A The dominant 55 kDa band represents the processed MPO heavy chain (amino acid residues 279-745). Two patient 8 samples have a strong 20 kDa band, a fragment representing a C-terminal MPO fragment (the antibody is specific for a peptide sequence near the C-terminus). The protein loading is not normalized for uromodulin content. Therefore, staining intensities do not allow direct quantitative comparisons. M_r_ standards (St) in gels from top to bottom are: 100, 75, 50, 37, 25, and 20 kDa.

Comparing datasets among the individual patients, we identified statistically significant variances in abundance for many proteins with functions in innate immunity (Figure S7, Suppl. Data). MPO and LTF data are depicted in Fig. 7A. MPO is the main ROS-generating enzyme in neutrophils. LTF is a multifunctional protein including an iron-sequestering function. Correlation analyses for numerous neutrophil and eosinophil effectors are displayed in Fig. 7B. This data shows that subsets of proteins cluster based on known enrichments in a specific type of cell or subcellular organelle: for instance, the eosinophil granule proteins bone marrow proteoglycan PRG2 and EPX and, as reported in (35), azurophilic granule proteins released by neutrophils: MPO, cathepsin G, azurocidin, and elastase. Such correlation data support the notion that the effectors contribute to immune defenses triggered by persistent biofilm formation on catheters. Western blots for MPO, not normalized for total protein in UP samples, confirm activation of neutrophils in patients in response to CB formation. MPO blots for selected timepoints pertaining to three patients, shown in Fig. 7C (P2-6, P7-55, P4-47), revealed low band intensities. This is consistent with low UP/CB biomasses for these timepoints (Fig. 3). Regardless of patient origin, neutrophils infiltrate the urothelium and release their effectors at the catheter surfaces when they harbor microbial biofilms.

### Antibiotic drug treatments influence composition and resilience of CBs, with instances of transient and permanent changes

A wound infection of patient P7 was treated with Bactrim, a combination of sulfamethoxazol and trimethoprim (TMT), at timepoint 41UP/CB. This treatment was stopped a week prior to timepoint 56UP/CB, resulting in the transient bacterial elimination at timepoint UP/CB55. *P. mirabilis* rapidly recolonized as shown in Fig. 3. A cohabitant of the P7 CBs, *H. influenzae*, also recurred as a minor component at post-treatment timepoints. The virulence factor HpmA was highly abundant in the *P. mirabilis* proteome prior to and after treatment, supporting recurrence of colonization with the same *P. mirabilis* strain. Disk diffusion DSTs for a bacterial isolate (CB65) from P7 revealed that it was susceptible to a TMT/sulfonamide combination (Table 2). Treatment of P9 with levofloxacin over ten days resulted in reduced microbial biomass at timepoint UP/CB75 and eliminated *S. marcescens* from the biofilm (Fig. 3). But *E. faecalis* and *P. aeruginosa* strains persisted, and *S. aureus* and *C. albicans* emerged as new CB community members at post-treatment timepoints. Fungal pathogens are not inhibited by fluoroquinolone drugs. We also isolated a slowly growing *E. faecalis* strain on blood and Mueller-Hinton agar from 74CB. This isolate was indeed resistant to a fluoroquinolone, ciprofloxacin (Table 2). *E. faecalis* has a high incidence of *parC* and *gyrA* mutations that confer fluoroquinolone resistance in strains causing UTI (36). We did not identify peptides displaying the common ParC and GyrA amino acid substitutions. A *S. marcescens* strain isolated from 74CB was susceptible to ciprofloxacin. We isolated a *S. aureus* strain from 75CB. This isolate was not only resistant to ciprofloxacin but also to ampicillin and cephaloxin, according to DSTs (Table 2). Several antibiotic drugs were administered to treat a P2 wound infection, with timepoints ranging from 10UP to 42CB (Fig. 3). Intravenous cephalosporin treatments eliminated strains of *S. aureus*, *E. faecalis*, *B. scardovii* and *P. aeruginosa* from the patient’s catheters, while *Brevundimonas* and *Bordetella hinzii* strains were more resilient. Daptomycin treatment diminished the *B. hinzii* colonization burden, from 32CB to 33CB, and *K. pneumoniae* emerged as the dominant species. We did not test the *K. pneumoniae* isolate from 43CB for resistance to daptomycin, a last-resort antibiotic to treat Gram-positive infections, but DSTs revealed the strain’s extended spectrum β-lactam and ciprofloxacin resistance (Table 2).

**Table 2.** Antimicrobial drug resistance of bacterial strains isolated from P1, P2, P7 and P9

| Bacterial species | Sample source | Ampicillin<br>100 µg | Cefotaxime<br>30 µg | Cephalexin<br>30 µg | Ciprofloxacin<br>5 µg | Sulfanilamide /<br>TMT 200µg/ 12µg | Gentamycin<br>30 µg |
| --- | --- | --- | --- | --- | --- | --- | --- |
| *P. aeruginosa* (P1) | 19CB | 0.6 | NP | NP | 2.7 | 0.6 | NP |
| *K. pneumoniae* (P2) | 43CB | 0.6 | 0.6 | NP | 0.6 | NP | 1.4 |
| *P. mirabilis* (P7) | 65CB | 2.2 | NP | 2.4 | 2.1 | 2.0 | 1.0 |
| *S. marcescens* (P9) | 74UP | 2.2 | NP | NP | 2.2 | 1.9 | NP |
| *E. faecalis* (P9) | 74UP | 2.0 | NP | 0.6 | 0.6 | 0.6 | NP |
| *S. aureus* (P9) | 75UP | 1.1 | NP | 0.8 | 0.6 | 3.2 | 1.6 |

All disk diffusion tests were performed in three replicates using Mueller-Hinton agar. The *E. faecalis* isolate was also grown on blood agar yielding the same results. The data provided here are growth clearance zones in cm (disk diameter 0.6 cm). Abbr: P: patient; NP: susceptibility tests were not performed. TMT: trimethoprim.

To gain further insights into antibiotic resistances of bacterial isolates observed in the DSTs, pan-proteome database searches including multiple genotypes for a species and ORFs derived from plasmids or pathogenicity islands were conducted. Thereby, database searches were expanded to “accessory proteomes”. Bacteria harvested from DST plates and clinical samples were analyzed. Proteins linked to various modes of antibiotic resistance are listed in Dataset S8 (Suppl. Data). The *S. aureus* isolate from 75UP (P9) expressed the penicillin-binding protein 2 MecA, β-lactamase BlaZ and aminoglycoside 4’-adenylyltransferase AadD. Expression of the proteins matched the strain’s reduced sensitivities to ampicillin, cephalexin and gentamycin, respectively. A *S. aureus* strain isolated from 53CB (P6), the patient diagnosed with a kidney infection, also expressed MecA. A *P. mirabilis*isolate from P6 (30CB) expressed aminoglycoside O-phosphotransferase (APH(3’)-Ia) and dihydrofolate reductase DfrA17, indicative of resistances to aminoglycosides and TMT, respectively. The *P. mirabilis* isolate from CB65 (P7) did not express these proteins. The *E. faecalis* isolate from 74UP expressed a dihydrofolate reductase (DfrE) and an aminoglycoside O-phosphotransferase (APH (3’)-IIIa). DfrE expression explains the observed TMT resistance (Table 2). In summary, some bacterial pan-proteome analyses corroborated evidence of single- or multi-drug resistant species co-colonizing patient’s catheters.

## Discussion

Our systems-level study on biofilms that grow on indwelling catheters of neurogenic bladder patients generates new insights into CB longitudinal dynamics. Meta-omics data demonstrate that dominant strains of distinct bacterial species recur in serially replaced catheters under challenge by the innate immune system and antibiotic drug treatments. Gaining these insights depends on clinical samples that CAUTI animal models cannot simulate. Such animal models cannot introduce the polymicrobial mixtures that human CBs are composed of. The range of micro- to anaerobic growth conditions that, we conclude, affect the CB profiles are impossible to mimic in CAUTI models. Culture-independent surveys identified many fastidious bacteria in CBs in the context of CAUTI and CAASB (22, 23, 37). We profiled strictly anaerobic bacteria cohabitating CBs in 6 of the 8 patients on the proteome level for the first time: *A. massiliense*, *P. lymphophilum*, *B. scardovii*, *V. parvula*, and *C. curvus*. Fastidious organisms more tolerant of oxygen (*A. urinae* and *G. sanguinis*) were identified as persistent colonizers of CBs. There is increasing awareness of the fact that such bacteria are more common causes of UTI and CAUTI in immunocompromised patients (38–40). The protection a biofilm offers makes it likely that such bacteria colonize hosts with functional immune systems. There is no evidence that the patients surveyed here, despite their spinal cord injuries, are immunocompromised. Animal models do not mimic the sequential replacement of catheters, a clinical need for many neurogenic bladder patients, because of the one-time catheter implantation technique. Low quantities of biological materials from murine catheters limit experiments requiring > 50 µL sample volumes (proteomics, metabolomics). Differences of murine *vs.* human urothelial surface markers (18), Toll-like receptors (41), cytokeratin abundance patterns in urothelial and stratified squamous epithelial cells (42, 43), and the lack of key innate immune effectors in murine neutrophils (e.g. defensin-1) encourage clinical investigations to study the crosstalk of polymicrobial biofilms with the host immune system and to conceptualize new ideas for therapeutic interventions.

In partial agreement with studies in which microbial colonization of recurrently catheterized patients was examined from UP samples (24, 25), we observed long-term bacteriuria for greater than 90% of the specimens. In agreement with another survey of longitudinally profiled catheter extracts (26), we observed frequent CB cohabitation by two or more pathogens and altered microbial community profiles following antibiotic drug treatments. *P. mirabilis, K. pneumoniae, P. aeruginosa, E. coli* and *E. faecalis* were most prevalent and persistent according to our data and (26). In contrast to the cited studies, we found fastidious bacteria to be common CB cohabitants derived from chronically catheterized patients. Spinal cord-injured cohorts, used in (25) and our study, are not associated with any specific subset of uropathogens. We cannot confirm previous reports where *P. stuartii* was found to be as common a cause of recurrent CAASB as *P. mirabilis* (24, 25). Our data confirm that *P. mirabilis* strains have significant fitness advantages, resulting in their persistence in and dominance of CBs, when catheters become encrusted with insoluble phosphate salts and are used by the bacteria to establish co-aggregates with salts (3, 11, 12, 26).

Our UP and CB proteome data strongly support the notion that distinct strains of a given microbe dominate the CB community of a given patient, as compared to a mixture of strains. While absolute evidence to back up this claim requires analysis of the strains’ genomes in a series of CBs, which we plan to do in future work, identification of genes not part of a species’ core genome in some but not other patients allow this conclusion. For instance, HpmA, a *P. mirabilis* cytotoxin, was identified only in P7 samples. Nrp (PMI2596-PMI2605), a *P. mirabilis* siderophore biosynthesis system, was expressed only in P1 and P4, but not in samples from P5, P6, and P7. An *E. coli* plasmid-encoded ferric iron uptake system (UTI89_P010-P017) and adhesion factor Hek were consistently observed in P1 samples but absent in *E. coli* proteomes present in the CBs of other patients. The *E. coli* aerobactin biosynthesis system Iuc and its receptor IutA were abundant in P1, P2 and P5, but not present in samples from P8 and P9. Subunits of the twitching mobility type IV pilus were identified in CBs that *P. aeruginosa* contributed to in P1, but not in P2, P8, and P9 samples. The type IV pilus is part of the *P. aeruginosa* accessory genome and itself facilitates conjugative DNA transfer (44). The *E. coli* aerobactin system and the adhesin Hek were reported to be expressed by virulent strains that cause urosepsis (45) and neonatal meningitis (46), respectively. Nrp enzymes synthesize a yersiniabactin-like siderophore. Few *P. mirabilis* genomes have been sequenced and annotated so that insights into Nrp gene cluster frequency for clinical strains other than HI4320 (47) are not yet available. Patient-specific identifications of ORFs considered part of accessory genomes argue in favor of a single dominant genotype (strain) for a species. But it cannot be ruled out that other strains make minor contributions to the same CBs. Dominant strains re-colonize catheters that are replaced in patients, either from intracellular or extracellular urinary tract reservoirs. Intra-urothelial bacterial communities have been described for *E. coli* and *K. pneumoniae* (48, 49). Our data clearly support re-colonization at the community level given that the same species recur in longitudinal CB series. That, in turn, points towards extracellular retention of polymicrobial clusters dispersed from catheter surfaces and persisting in the urinary tract via mucosal adherence. Important adhesion proteins of Gram-negative bacteria in the urinary tract are fimbriae which varied in abundance in our datasets. Highly abundant were the MR/K fimbriae of *K. pneumoniae* (KPN03276-KPN03280 (50)) and MR/P fimbriae of *P. mirabilis* (PMI0263-PMI0270 (51)). The MrpA variance in abundance across datasets is displayed in Fig. 4. Each species produced ECP-type fimbriae (KPN00290-KPN00295, PMI2997-3003), but in lower quantities. Fimbriae display phase variation and permit the rapid adaptation to conditions that foster resilience in the host milieu, including the attachment to biotic surfaces and medical devices (52, 53). Mannose-resistant MR/K and MR/P fimbriae, as well as the *P. aeruginosa* type IV pilus (54), are involved in cellular aggregation and biofilm formation. Consistent with our data and the literature, we conclude that these surface assemblies provide fitness advantages and mediate the persistence of strains that express them in the catheterized human urinary tract.

The proteomic data provided insights into the energy metabolism and transition metal acquisition of bacterial species simultaneously present in CBs derived from clinical samples, to our knowledge for the first time. The discussion focuses on *P. mirabilis, E. coli, and E. faecalis* because these bacteria formed mixed CB communities recurrently in several patients. Each species expressed at least two ABC transporters known or predicted to facilitate TMI import. The Gram-negative species expressed more than one biosynthesis system for siderophores and TonB-dependent receptors for their uptake when complexing Fe^3+^ and other TMIs. We assessed abundance correlations among the systems that facilitate TMI acquisition (Fig. 5). The correlations, derived from *P. mirabilis, E. coli*, and/or *E. faecalis* over a series of timepoints, were positive in CB profiles for P1, but not for P4 and P5. We hypothesize that these results reflect robust, simultaneous growth of the pathogens in P1. Growth requires TMI uptake in a metal ion-sequestering host milieu. TMIs are incorporated as cofactors or components of cofactors such as Fe-S and heme into enzymes that support metabolism in the cell, particularly energy metabolism. Several studies have established links between TMI acquisition in a TMI-starved host milieu and the expression levels of Fe and Fe-S cluster-harboring enzymes that support the bacterial energy metabolism (55, 56). Thus, we consider the observed correlation data supportive of active metabolism and growth in *P. mirabilis, E. coli*, and/or *E. faecalis* in P1. They may act cooperatively and competitively as it pertains to the uptake of TMIs (57). The cellular energy going into the assembly of the TMI acquisition systems in the host environment appears to be high. For example, the *E. coli* strain in P1 highly expressed systems for aerobactin, yersiniabactin and enterobactin synthesis and uptake as well as heme uptake. Some of their components correlated with the abundance of the only characterized *E. faecalis* system for TMI acquisition, EfaABC. This ABC transporter is regulated by Mn^2+^ (58) and promotes bacterial growth under Mn^2+^ starvation conditions (32). Whether EfaABC can capture Mn^2+^ or other trace metals from *E. coli* or *P. mirabilis* siderophores or heme complexes remains to be shown. We expressed EfaA and another *E. faecalis* lipoprotein predicted to be part of an ABC transport system for TMIs (EF0577) recombinantly. Both proteins revealed biochemical evidence of Co^2+^ binding *in vitro*. An EfaA ortholog, PsaA, binds Mn^2+^ and Zn^2+^ ions but only transports Mn^2+^ into the *Streptococcus pneumoniae cell* (59). We hypothesize that the growth states of *P. mirabilis, E. coli*, and/or *E. faecalis* in P4 and P5 were variable, perhaps including different levels of quiescence, a known trait of persister cells in biofilms (60).

Furthermore, we hypothesize that communications among cohabitating bacteria affecting TMI homeostasis occur, a process that has been studied using model systems (57, 61). The human immune defense system perturbs this homeostasis. We surveyed proteins involved in sequestration of iron (LTF), zinc (calgranulins), and enterobactin (lipocalin-2). Like antimicrobial effectors such as the ROS-generating enzymes MPO and defensin-1, the proteins are released by neutrophils (or eosinophils) and abundant in human proteomes of UP and CB samples (Table 1). TMI-sequestering proteins starve bacteria of these essential cofactors at the CB-urothelial interface. At least three Gram-negative species profiled in CBs expressed high quantities of biosynthetic systems to produce lipocalin-2-insensitive siderophores: *P. aeruginosa* pyoverdin in P1, *E. coli* aerobactin in P1, P2 and P5, *E. coli* yersiniabactin in P1, P2, and P8; a *P. mirabilis* yersiniabactin-like siderophore in P1 and P4. Lipocalin-2 insensitivity likely increases the resilience of strains residing in CBs. Siderophore receptors operate in tandem with a TonB-dependent energy-transducing system (TonB-ExbBD), which provides the energy for the transport of TMIs across the outer membrane. A single TonB energy-transducing system seems to be expressed by most Gram-negative bacteria, and the system’s structure has been determined (62). TonB-ExbBD is an important molecular target to discover or design inhibitors. The system may disable TMI uptake in the human host when infections occur. Chemical compounds were screened to identify inhibitors of siderophore uptake using *E. coli* and *A. baumannii* TonB strains (63, 64). Such inhibitors have the potential to strengthen nutritional immunity and may be non-toxic in mammals given that TonB-ExbBD structures are unique to Gram-negative bacteria.

Finally, we observed different outcomes in three cases of systemic antibiotic treatments. Of note, the antibiotic drugs did not target microbial pathogens present in CBs, but those suspected to cause a comorbidity, chronic wounds. In P7, the *P. mirabilis* biofilm disappeared during Bactrim intake, but the strain regrew when antibiotic drug treatment was arrested. A CB isolate from this patient was sensitive to TMT/sulfanilamide treatment. In P9, a biofilm consisting of *E. faecalis, P. aeruginosa*, and *S. marcescens* was exposed to LEV treatment. This resulted in *S. marcescens* elimination from the biofilm. The other species persisted. *S. aureus* and *C. albicans* joined the reestablished biofilm. *E. faecalis* and *S. aureus* isolates from relevant CB timepoints were resistant to the fluoroquinolone CIP. P2 was treated sequentially with three different drugs over six weeks. Most pathogens were eliminated, but treatments failed to prevent sequential outgrowth of species not susceptible to the administered drugs, first *B. hinzii* (cephalosporin treatment), then an ESBL-resistant *K. pneumoniae* strain and *C. albicans* (daptomycin and fluoconazole treatments). Using pan-proteome searches for bacterial isolates grown *in vitro*, we identified proteins that cause antibiotic drug resistances in UTI and CAUTI pathogens, a serious public health problem threatening to lead to untreatable infections in the coming decades (65). The *S. aureus* strain from 75CB expressed BlaZ and MecA, proteins induced in expression via cross-talk among regulators of the corresponding genes (66). Additionally, the anoxic environment in deeper layers of biofilms and quiescence allow bacteria to survive during drug treatments even if the genomes do not harbor specific antibiotic resistance genes or drug efflux pumps (60). CAASB in chronically catheterized patients is a difficult medical problem. This study highlights the microbial escape routes and adaptability of polymicrobial communities formed on a medical device in the human body, following exposure to antibiotic drugs, nutritional starvation (e.g. oxygen and TMIs) and antimicrobial effectors of the immune system.

## Methods

### Ethics Statement

The Southwest Regional Wound Care Center (SRWCC) in Lubbock, Texas, and the J. Craig Venter Institute (JCVI, Rockville, Maryland) created a human subject protocol and a study consent form (#56-RW-022), which were approved by the Western Institutional Review Board (WIRB) in Olympia, Washington, followed by JCVI’s IRB in 2013. All human subjects were adults and provided written consent. The specimens were collected firsthand for the purpose of this study. There was a medical need to serially replace indwelling Foley catheters in patients due to their affliction with neurogenic bladder syndrome. Scientists at the JCVI did not have access to data allowing patient identification. Medical metadata for the subjects were encrypted. Electronic and printed medical records at the clinical site were retained for 4 years to facilitate the integration of medical and molecular research data.

### Human subjects and study design

Nine human subjects enrolled in this prospective study. They had irreversible spinal cord injuries (SCIs) and suffered from neurogenic bladder syndrome. Catheter replacement was part of routine patient care at SRWCC. Medical data included gender, ethnicity, antibiotic use, diagnosis of chronic wound infections, and diabetes. Facilitated by medical staff, the enrolled subjects provided 3 to 15 specimens (urethral catheters and urine from catheter collection bags) collected longitudinally at the clinic in 1- to 4-week intervals, depending on the number of visits that patients and physician agreed upon. Catheters were cut into 1-inch pieces, placed in polypropylene tubes and stored at −20°C, minimizing external contamination by use of gloves and sterile razor blades. Urine samples were obtained from catheter bag ports swabbed clean with alcohol prior to collection. Urine aliquots of 20 to 50 ml were stored at −20°C. Infrequent draining of catheter bags may have allowed some *ex vivo* microbial growth in urine specimens. We assume that the quantitative ratios of microbes in urine sediments on the collection dates do not reflect those in the urine excreted over time. Containers in which specimens were stored were kept frozen during transport and transferred to a −80°C freezer until further use at JCVI.

### Urine and catheter specimen extraction and protein solubilization for proteomics

The catheter materials were latex in the case of 8 patients and silicone (patient P7). The pH, color and turbidity of urine specimens and the crystallization of salts inside and on the surface of catheter segments were noted. To obtain a urine pellet (UP) from a sample, an aliquot was thawed, adjusted to 20°C and, if acidic, neutralized with 1 M Tris-HCl (pH 8.1) to a pH of ~ 6.5 to 7.5, and centrifuged at 3,200 × g for 15 minutes. UPs were aliquoted for rDNA and protein extractions and spare samples in ratios of approximately 10%, 45% and 45%, respectively. Urethral catheter pieces were extracted in two steps. Submerged in 100 mM sodium acetate (pH 5.5), 20 mM sodium meta-periodate, and 300 mM NaCl, catheter pieces were agitated in an ultrasonic water bath for 10 min at 20°C. The supernatant was recovered and concentrated. This was followed by two solubilization cycles of the pellet with a denaturing SED solution (1% SDS (v/v), 5 mM EDTA and 50 mM DTT) including 3 min heat treatment at 95°C. Two supernatants were recovered that we termed CB-1 (Na-acetate buffer) and CB-2 (SED solution). The extraction of UP samples was limited to solubilization in SED solution. Experimental details were described previously (28, 31). All centrifugal centrifugation steps were performed using Ultrafree-4 filter units (10 kDa MWCO), potentially eliminating small peptides from the concentrates. However, we determined that peptides with M_r_ values lower than 5 kDa (e.g. neutrophil defensin-1; 3-4 kDa) were partially retained. Processing steps for the solubilized UP and CB samples were based on the Filter-Aided Sample Preparation (FASP) method (67), adapted by us to the use of 100 µg protein for digestion with sequencing-grade trypsin in 50:1 ratios in Vivacon 10k filters (Sartorius AG, Germany) (28). FASP-processed peptide mixtures were desalted using the Stage-Tip method (68) and lyophilized for LC-MS/MS proteomic analysis.

### Shotgun proteomics via LC-MS/MS

Desalted peptide mixtures derived from UP, CB-1, and CB-2 samples were dissolved in 10 μl 0.1% formic acid (solvent A) and analyzed using one of two LC-MS/MS systems: (1) a high-resolution Q-Exactive mass spectrometer (MS) coupled to an Ultimate 3000-nano LC system; (2) a low-resolution LTQ-Velos Pro ion-trap mass spectrometer coupled to an Easy-nLC II system. Both systems (Thermo Scientific, San Jose, CA) were equipped with a FLEX nano-electrospray ion source at the LC-MS interface. Analytic procedures were previously described for the Q-Exactive (69, 70) and LTQ-Velos Pro (71) platforms. For LTQ-Velos Pro analysis, peptides present in a sample were trapped on a C_18_ trap column (100 μm × 2 cm, 5 μm pore size, 120 Å) and separated on a PicoFrit C_18_ analytical column (75 μm × 15 cm, 3 μm pore size, 150 Å) at a flow rate of 200 nl/min. Starting with solvent A, a linear gradient from 10% to 30% solvent B (0.1% formic acid in acetonitrile) over 195 minutes was followed by a linear gradient from 30% to 80% solvent B over 20 min and re-equilibration with solvent A for 5 min. Following each sample, the columns were washed thrice using a 30-min solvent A to B linear gradient elution to avoid sample carry-over. For Q-Exactive analysis, LC was conducted as reported earlier (69). Electrospray ionization was achieved by applying 2.0 kV distally via a liquid junction. Parallel to LC gradient elution, peptide ions were analyzed in a MS^1^ data-dependent mode to select ions for MS^2^ scans using the software application XCalibur *v2.2* (Thermo Scientific). The fragmentation modes were collision-induced dissociation (CID) with a normalized collision energy of 35% (LTQ-Velos Pro) and higher-energy collisional dissociation (HCD) with a normalized collision energy of 27% (Q-Exactive). Dynamic exclusion was enabled. MS^2^ ion scans for the same MS^1^ *m/z* value were repeated once and then excluded from further analysis for 30s. Survey (MS^1^) scans ranged from a *m/z* range of 380 to 1,800 followed by MS^2^ scans for selected precursor ions. Survey scans with the Q-Exactive were acquired at a resolution of 70,000 (m/Δm) with a *m/z* range from 250 to 1,800. MS^2^ scans were performed at a resolution of 17,500. The ten most intense ions were fragmented in each cycle. Ions that were unassigned or had a charge of +1 were rejected from further analysis. Two or three technical LC-MS/MS replicates were run for UP, CB-1 and CB-2 peptide extracts.

### Computational proteomic data analyses and proteome quantifications

The raw MS files were combined for database searches as follows: (1) all technical replicates for a given UP sample; (2) all replicates from both CB-1 and CB-2 fractions for a given CB sample. The Sequest HT algorithm integrated in the software tool Proteome Discoverer v1.4 (Thermo Scientific) was used as the search engine with analytical parameters described previously (69, 70). Only rank-1 peptides with a length of at least seven amino acids were considered. The FDR rates were estimated using the integrated Percolator tool with a (reverse sequence) decoy database. Protein hits identified with a 1% FDR threshold were accepted for data interpretation. The ‘protein grouping’ function was enabled to ensure that only one protein was reported when multiple proteins shared a set of identified peptides. The database contents were the reviewed protein sequence entries in the non-redundant Human UniProt database (release 2015-16; 20,195 sequences) and sequence entries for 23 microbial genomes reported as major causes of UTI (69). Based on 16S rRNA genus assignments and iterative proteomic searches using database subsets for distinct species part of the same genus, the proteomic searches were customized for the same series from each patient. Iterative searches followed by database customization were also applied to samples containing one or potentially more members of the Enterobacteriaceae family. The entire list of species searched in this process is listed in Table S3 (Suppl. Data). The pan-proteome database searches for five distinct microbial species were performed by downloading non-redundant pan-proteome sequence databases from UniProt-Proteome using the Sequest HT algorithm as mentioned above. MS raw files were deposited in PRIDE (via ProteomeXchange) with the identifier PXD012048. The sums of peptide-spectral match counts (PSMs) assigned to each microbial species and *Homo sapiens* were the basis for the individual species-based quantification in both UP and CB samples. Relative to the total proteome (all PSMs) in a dataset, normalized abundance values were obtained for all species in the Proteome Discoverer v1.4-derived dataset. They were the basis of quantitative displays in Fig. 3 and Figure S5 (Suppl. Data). For quantitative analyses that assessed protein variances, of human and microbial species origins, normalization was based on total PSMs of the respective species. This normalization pertains to proteomic data displayed in Figs. 4, 5, 6 and 7, Table 1, Dataset S6 and Figure S7.

### Microbiota analyses

Microbial cell lysis and DNA extraction methods from catheter extracts and UP samples (31) and the amplification of V1-V3 regions of the 16S rDNA bacterial genes as well as sequencing on the MiSeq sequencer from Illumina Inc. were described previously (72). The UPARSE pipeline for the phylogenetic analysis was used (73). OTUs were generated *de novo* from raw sequence reads using default parameters in UPARSE, the Wang classifier and bootstrapping using 100 iterations. The taxonomies were assigned to the OTUs using Mothur and applying the SILVA 16S rRNA database version 123 as reference database (74). Unbiased, metadata-independent filtering was applied at each level of the taxonomy by eliminating samples with less than 2000 reads and OTUs present in less than ten samples. Filtered data were analyzed based on the relative contributions of microbial genera in a distinct sample. The Shannon index was used to measure the alpha diversity.

### Microbial cultures

Surviving bacteria present in frozen catheter extracts and UP samples stored at −80°C were cultured on BHI broth, blood and MacConkey agar plates to isolate strains. Single colonies were picked, Gram-stained and microscopically assessed before culture stocks were generated either directly from a plate or a 5-10 mL overnight BHI suspension culture. Stocks were frozen in 10% glycerol at −80°C. To determine the identity of bacterial species, cell lysis, proteomic sample processing and database search methods we mentioned in the context of clinical sample analysis were used. If a dataset suggested impurities, bacterial strains were streaked out again from the original stock and new colonies were picked and reanalyzed. The OD_600_ values reached for *E. faecalis* strains in BHI media were below 0.6, the values for species of the Enterobacteriaceae family were usually higher than 0.8. Some fastidious bacterial species were isolated and lysed directly from blood agar plates for proteomic analysis. The isolation of other fastidious bacteria derived from anaerobic storage in liquid nitrogen and anaerobic culture, without freezing catheter specimens, was previously reported (31). Further details are provided in Dataset S2 (Suppl. Data).

### Expression vectors, gene cloning, and recombinant protein expression

Two expression vectors were used to clone EF2076, EF0577 and EF3082 (*E. faecalis* V583) into the *E. coli* strain BL21(DE3)/pMagic (75). The ccdB cassette was inserted into the cloning region to enhance the efficiency of screening for the correct clone. pMCSG53 encodes the N-terminal His-tag (75). PCR primers in Table 3 were used to amplify the ORFs excluding export signal sequences. Inserted DNA fragments were prepared using standard PCR protocols using Phusion polymerase (New England Biolabs). The pMCSG53 vector was linearized by cleavage of the cloning site with SspI, and PCR products were cloned using the Gibson Assembly method (76) and transformed into *E. coli* DH5α. Sequences of the clones were validated by Sanger sequencing. Their transformation into BL21(DE3)/pMagic was followed by growth at 37 °C in 0.5 L LB containing 100 μg/ml ampicillin to an OD_600_ of ~ 0.8 when expression was induced by adding 1 mM IPTG. Following incubation at 20°C for 16-18 hours, the cells were lysed. *E. faecalis* target proteins were purified on Ni-NTA resin, cleaving the affinity tags with TEV protease as previously reported (77). Purity and solubility of the purified proteins was assessed by Nu-PAGE.

**Table 3.**
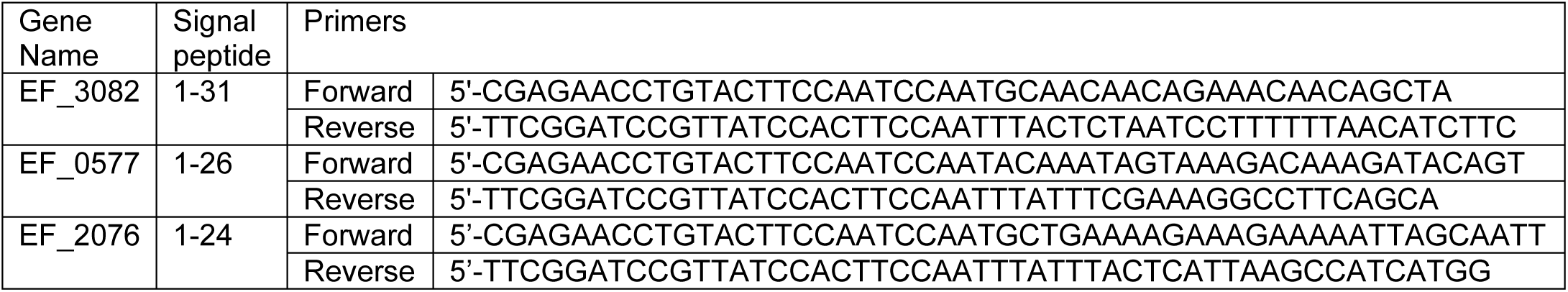
Sequences of PCR primers

### Thermal shift assay

Interaction of proteins with a small molecule can increase its stability and the melting temperature (T_m_) (78). To denature each of the purified *E. faecalis* proteins, a 1 mg/ml solution was incubated with 1 ml 6 M guanidine-HCl (pH 8). To renature the protein, 50 μl Ni-NTA agarose resin was added. The suspension was incubated gently agitating at 4°C for 1 h. The flow-through fraction was discarded. The resin was washed with 8 M urea/PBS, and the protein gradually renatured via buffer exchange into PBS followed by elution in 50 μl of an elution buffer containing 250 mM imidazole. Renatured proteins were either left in PBS or CoCl_2_ was added at a final concentration of 50 μM. Ten μl of a protein preparation (0.5 mg/ml) was placed in a 384-well plate adding 2 μl 100X SYPRO Orange. Temperature melting profiles were monitored in a Light Cycler 480 (Roche Life Science).

### Disk diffusion antibiotic drug susceptibility tests

We followed guidelines from a Manual of Antimicrobial Susceptibility Testing (79) to perform drug susceptibility tests via disk diffusion. All antibiotics were dissolved as specified by the Korean society of laboratory medicine manual and placed on 0.6 mm disks in concentrations as follows: Ampicillin (100 µg); Cefotaxime (30 µg); Cephalexin (30 µg); Ciprofloxacin (5 µg); Sulfanilamide/TMT (200µg/ 12µg); and Gentamycin (30 µg). The McFarland standard (0.5) was used to adjust bacterial turbidity and plate approximately 1.5 × 10^8^ CFU/ml of bacterial inoculum from a ~ 6h BHI culture (Gram-negative) or a Mueller-Hinton agar plate (Gram-positive). Clearance zones were measured after 16 hours of growth at 37°C in a 5% CO_2_ incubator.

### Statistical analysis methods

To determine statistical changes in taxonomic α-diversity, Wilcoxon rank sum tests were performed. These tests were also used to assess variation in the abundances of proteins in proteomic datasets from different patients. Analysis of variance (ANOVA) was used to compare the variation between patients. We applied a P-value cutoff of 0.05 to determine statistically significance. The correlation plot was derived from the corrplot package in R, using the Pearson correlation with a P-value less than 0.05. Box plots with statistically significance data were generated using ggsignify and ggplots in R.

## Acknowledgements

This work was supported by the National Institutes of Health grant R01GM103598 titled “Urethral catheter-associated polybacterial biofilm formation and dispersal”. The funder had no role in study design, data collection and interpretation, or decisions to submit the work for publication. We thank the Ruggles Family Foundation for the support in acquiring the Q-Exactive mass spectrometer.

